# REDUCED FOLATE CARRIER 1 IS PRESENT IN RETINAL MICROVESSELS AND CRUCIAL FOR THE INNER BLOOD RETINAL BARRIER INTEGRITY

**DOI:** 10.1101/2022.10.14.511731

**Authors:** Gokce Gurler, Nevin Belder, Mustafa Caglar Beker, Melike Sever-Bahcekapili, Gokhan Uruk, Ertugrul Kilic, Muge Yemisci

## Abstract

**Background:** Reduced folate carrier 1 (RFC1; SLC19a1) is the main responsible transporter for the B9 family of vitamins named folates, which are essential for normal tissue growth and development. While folate deficiency resulted in retinal vasculopathy, the expression and the role of RFC1 in blood-retinal barrier (BRB) are not well known.

**Methods:** We used whole mount retinas and trypsin digested microvessel samples of adult mice. To knockdown RFC1, we delivered RFC1-targeted short interfering RNA (RFC1-siRNA) intravitreally; while, to upregulate RFC1 we delivered lentiviral vector overexpressing RFC1. Retinal ischemia was induced 1-hour by applying FeCl_3_ to central retinal artery. We used RT-qPCR and Western blotting to determine RFC1. Endothelium (CD31), pericytes (PDGFR-beta, CD13, NG2), tight-junctions (Occludin, Claudin-5 and ZO-1), main basal membrane protein (Collagen-4), endogenous IgG and RFC1 were determined immunohistochemically.

**Results:** Our analyses on whole mount retinas and trypsin digested microvessel samples of adult mice revealed the presence of RFC1 in the inner BRB and colocalization with endothelial cells and pericytes. Knocking down RFC1 expression via siRNA delivery resulted in the disintegration of tight junction proteins and collagen-4 in twenty-four hours, which was accompanied by significant endogenous IgG extravasation. This indicated the impairment of BRB integrity after an abrupt RFC1 decrease. Furthermore, lentiviral vector-mediated RFC1 overexpression resulted in increased tight junction proteins and collagen-4, confirming the structural role of RFC1 in the inner BRB. Acute retinal ischemia decreased collagen-4 and occludin levels and led to an increase in RFC1. Besides, the pre-ischemic overexpression of RFC1 partially rescued collagen-4 and occludin levels which would be decreased after ischemia.

**Conclusion:** In conclusion, our study clarifies the presence of RFC1 protein in the inner BRB, which has recently been defined as hypoxia–immune-related gene in other tissues and offers a novel perspective of retinal RFC1. Hence, other than being a folate carrier, RFC1 is an acute regulator of the inner BRB in healthy and ischemic retinas.

## BACKGROUND

The Reduced Folate Carrier 1 (RFC1), also called solute carrier family 19 member 1 (SLC19A1/SLC19a1) is responsible for transporting folates via Blood Brain Barrier (BBB) to the brain parenchyma (1, 2). Folates belong to the B9 family of vitamins and are essential for proper functioning of the Central Nervous System (CNS) as they are crucial for many key processes such as histone and DNA methylation, DNA replication and repair, RNA synthesis, methionine production, homocysteine remethylation, aminoacid metabolism, neurotransmitter, and phospholipid synthesis (3–6). Several clinical studies in adults determined that lower serum folate levels could cause neurological dysfunction (7, 8). Moreover, folate deficiency was shown to result in microvascular complications in the retina (9, 10). However, the findings of these studies are chronic consequences that occur with long-term changes and/or deficiency of folates.

Retina is the extension of the CNS, displaying similarities in terms of anatomy, function, and response to insults (11). The retina has the blood-retina barrier (BRB) that precludes the free transfer of substances from the capillary blood, and is composed of two parts (12). The outer BRB is constituted of the retinal pigmented epithelium (RPE), which are specialized epithelial cells sealed with tight junctions, residing between the neural retina and a fenestrated capillary network called choriocapillaris. It partly functions to determine the transfer of nutrients from the choriocapillaris to the retina (13). The inner BRB is established in the inner retinal microvessels and comprises specialized endothelial cells secured by tight junctions, enclosed by basal lamina, pericytes, and perivascular astrocytes; hence it is essential for the maintenance of homeostasis through its selective properties such as harboring specialized nutrient carriers (12). To date the expression of RFC1 and its transporter function in the retina was primarily evaluated only in the RPE, thus the outer BRB (14–17).

The recent cerebral microvessel transcriptome data showed prominent RFC1 presence in pericytes, the cells crucial with various roles in the regulation of microvascular blood flow and barrier integrity. The retinal microvessels have similar features to the brain (18–20) and are unique for having the highest pericyte ratio in the body. Therefore, our first aim was to investigate the presence of RFC1 protein in the inner BRB, and especially in the pericytes and endothelial cells.

RFC1 has been recently shown in the cell lines to be an importer of immunotransmitter cGAMP and other cyclic dinucleotides with robust antitumoral immune response (21) and considered to be the hypoxia–immune-related gene in multiple myeloma patients (22). The expression and activity of RFC1 was shown to differ during insults like hyperglycemia, nitric oxide exposure, hyperhomocysteinemia and oxidative stress in the RPE (14, 15, 17). However, no *in vivo* studies were carried out in the inner BRB. Therefore, our second aim was to investigate the role of RFC1 protein in the inner BRB in healthy retinas and in acute retinal ischemia which is a world-wide problem for blindness.

In this study, we elucidated the presence of RFC1 protein in the endothelial cells and pericytes of the inner BRB. We employed a custom-designed RFC1-targeted Accell short interfering RNA (siRNA) to silence the RFC1 gene, and a lentiviral vector to overexpress RFC1 in the retinas *in vivo*. Abrupt and brief changes in RFC1 caused a disruption in the barrier properties of the retinal microvessels. Thus, we identified the immediate role of RFC1 protein in maintaining the inner BRB in health and in acute retinal ischemia.

## METHODS

### Animals

Experiments were performed according to the approval provided by Hacettepe University Animal Experimentations Local Ethics Board (No: 2019/13-06 and 2021/03-19). Adult female and male Swiss albino mice (25–35 g) housed under 12 hours of light and 12 hours of darkness at standard room temperature (22±2°C) and 50-60% humidity was used for *in vivo* experiments. Mice were allowed water and food ad libitum. The recommendations of ARRIVE guidelines and institutional instructions were considered in housing, care, and while performing experiments, the experimental groups were decided in a randomized and blinded manner. For all the surgical procedures intraocular injections, the animals were anesthetized with ketamine (80 mg/kg, intraperitoneal injection, Ketalar®, Pfizer, Cambridge, United Kingdom) and xylazine (8 mg/kg, intraperitoneal injection, Alfazyne® 2%, Alfasan International, Woerden, the Netherlands). A rectal probe connected to the homeothermic blanket with a control unit (Harvard Apparatus, U.S.A.) was utilized to control the body temperature (37.0 ± 0.5°C). Pulse rate and oxygen saturation were monitored from the right lower limb via an oximeter (The LifeSense® VET Pulse Oximeter, Nonin Medical Inc., USA). Vital signs (rectal temperature, blood pressure, oxygen saturation, and pulse rate) were kept within physiological limits and recorded. After operations, mice were kept on a homoeothermic blanket until they recovered from anesthesia fully. Next, the mice were kept individually in their cages until sacrificed.

### Retinal Whole Mount Preparation

The eyeballs harvested were immersed in 4% paraformaldehyde (PFA) for 1 hr, and under a stereomicroscope, we made a puncture at the corneal border with an insulin needle while eyeballs were resting in one drop of PBS at room temperature. A circular cut along the limbus was made with the help of a micro-dissector. Following the extraction of the lens and vitreous body, the intact retina was detached from the sclera via a 45° thick curved tip tweezer by going around the edges. Then, the flattened retina was placed into a centrifuge tube consisting of 200 μL PBS for immunohistochemistry.

### Retinal Coronal Section Preparations

Obtained eyeballs were immersed in 4% paraformaldehyde (PFA) for 24 hr followed by incubation in 30% sucrose for 2 days. Next, 20 μm sections were obtained by cryostat.

### Retinal Trypsin Digest Preparation

We used modified retinal trypsin digestion protocol to isolate retinal microvessels from the surrounding tissue (23). Retinas were separated in cold Phosphate Buffered Saline (PBS), then immersed in 250 μL filtered ddH_2_0, and, at room temperature, kept on a shaker until they disintegrated. Then kept in 0.3% trypsin (PAN Biotech 1:250) 0.1 M Tris buffer (pH 7.8) in the incubator adjusted to 37°C. The separation of microvessels were seen clearly, tissue was washed with dH_2_O, and the procedure was repeated until no debris remained. Isolated retinal microvessels were transferred to poly-L-lysine coated slides without disrupting their integrity, and air-dried. This method allowed isolation of the retinal vessels containing endothelial cells, basal lamina, pericytes, the attached perivascular astrocyte processes, without the presence of other cell types like photoreceptor cells.

### Immunohistochemical Studies

For immunohistochemical studies whole mount retinas, coronal sections and isolated retinal microvessels were used. PFA fixed whole mount retinas were kept in in 0.5% PBS-Triton-X, freezed 15 min at −80°C and thawed for 15 min at room temperature following overnight incubation in 2% TritonX-100 (Merck Millipore, 1086031000) PBS for permeabilization. They were immersed in 10% normal goat serum for blocking steps for 2 hr at room temperature or overnight at +4°C. Next, whole mount retinas were incubated at 4°C with primary antibodies against RFC1 (SLC19a1, MyBioSource MBS9134642 or Sigma-Aldrich AV44167) for three consecutive overnights. We used two commercial polyclonal antibodies developed against different non-overlapping epitopes of RFC1. RFC1/SLC19a1 antibody from MyBioSource (MBS9134642) was produced against a recombinant fusion protein containing a sequence related to amino acids 452-591 of human SLC19A1 (NP_919231.1), while RFC1/SLC19a1antibody from Sigma-Aldrich (AV44167) was produced against Synthetic peptide directed towards the N terminal region of human SLC19A1 (NP_919231). Then retinas were subsequently incubated with ‘Fluorescein’ or ‘Texas Red’ labeled Lectin (Vector Laboratories, Burlingame, CA). Retinal microvessels obtained via retinal trypsin digestion were permeabilized with 0.3% TBS-Triton-X for 30 min, blocked in 10% normal serum of the host of the associated secondary antibody for 1 hr at room temperature. Vessel webs were incubated at +4°C overnight with primary antibodies against RFC1; for mural cells against PDGFR-β (R&D Systems, AF1042), NG2 (Sigma AB5320A4), CD13 (Acris Antibodies, AM26636AF-N), for endothelial CD31 (BD Bioscience, 550274); for tight junctions ZO-1 (Sigma, MAB1520), Claudin-5 (Invitrogen™, 35-2500), and for Collagen-4 (Abcam, ab6586) (n=3-6/marker). After several washes, the vessel webs were incubated for 1 hr at room temperature with appropriate Cy2-, Cy3- or Cy5-conjugated anti-rabbit IgG (Jackson ImmunoResearch 111-165-144 or 711-175-152) or Cy3-conjugated anti-rat IgG (Thermo Scientific™, A10522, 1:200). To assess the functional inner BRB integrity, PFA fixed whole mount retinas were incubated with Cy3-conjugated anti-mouse IgG (Jackson ImmunoResearch, 115-165-003) at +4°C overnight. Later, retinas were incubated with ‘Fluorescein’ or ‘Texas Red’ labeled Lectin (Vector Laboratories, Burlingame, CA) at +4°C overnight to visualize the vessels. Washed with TBS for 3 times, 5 min each; retinal tissues or retinal microvessels were mounted with a PBS/Glycerol mounting medium consisting of the nuclear staining Hoechst 33258 (Invitrogen™, 1:1000 dilution).

### Western Blotting

Whole retinas were lysed in Radioimmunoprecipitation assay (RIPA) buffer, sonicated, and homogenized on ice, then centrifuged at 10,000xg at +4°C for 20 min. The concentration of the lysates was assessed using BCA protein assay kit (Thermo Scientific™, 23225). After loading proteins (35 μg/well) to a homemade SDS-PAGE gel, 120 Voltage were applied for 1.5 hr. Proteins were then transferred to PVDF membranes by applying 120 mAmp/membrane for 2 hr at room temperature. Membranes were incubated with 5% BSA in 0.1% TBS-Tween for 1 hr at room temperature for blocking step and incubated with 1:1000 anti-RFC1 antibody produced in rabbit (Sigma-Aldrich; AV44167) at +4°C overnight. After washing with 0.1% TBS-Tween, we incubated the membranes with 1:5000 goat anti-rabbit HRP conjugated Invitrogen™, 31460) solution for 2 hr at room temperature. For detection, the membranes were treated with a high sensitivity enhanced chemiluminescent (ECL) substrate (Thermo Scientific™, 34096). Imaging was done by Kodak 4000M Image Station. β-Actin and β-tubulin III (Chemicon, MAB1637) was employed as an internal standard for cell culture homogenates and tissue homogenates respectively. Densitometric measurements were done by ImageJ 1.52 version (NIH, Bethesda, Maryland), and expressed as ratios to the internal standard.

### Imaging and Analysis

The images of the stained whole mount retinas and retinal microvessels were obtained with a Leica TCS SP8 confocal laser scanning microscope (Leica, Wetzlar, Germany) with a diode laser 405, 638 and OPSL 488, 552 nm, with a X, Y, and Z-movement controller, and a high-resolution PMT (Zeiss, Oberkochen, Germany) and HyD (Leica) detectors. Images were either acquired in a single focal plane while keeping settings constant between experimental or control groups for analysis or Z-stack mode with 0.80 μm wide steps along the Z axis. For analysis of immunostained retinal microvessels, 1024×1024 pixels and 276.79×276.79 μm sized, middle of the Z axis level images was used. Acquired images were exported as 8-bit grayscale .tiff formats from Leica Application Suite X (LAS X; version 3.5.5.19976) and opened in ImageJ 1.52 version. First, the lectin frame of each image was filtered by Gaussian blur (sigma=4) to reduce the random noise, and Huang Thresholding was applied to mask the lectin-positive microvessel areas of the images. After de-speckling (Process tab>Noise>Despeckle) and eroding (Process tab>Binary>Erode), the binary image was divided by 255 using Math function (Process tab>Math>Divide) to convert microvessel pixels to “1” and background pixels to “0”. Later, either “RAW” lectin image or the overlapping grayscale image belonging to tight junctions or collagen-4 was divided by the binary of the lectin image. This created 32-bit grayscale images in which microvessels had singular intensity and the background had infinity. Finally, the mean grey value (RFU, range between 0-255) was obtained which corresponded to the mean intensity of the pixels occupying the microvessel area. Number of pericytes were quantified from Lectin and Hoechst labelled microvessel preparations by researchers blinded to the experimental or control groups. As pericytes are well known to have protruding nuclei, they were easy to distinguish morphologically and spatially from endothelial cells having flat nuclei. The total microvessel length was measured with AngioTool (v 0.6a (64 bits), October 2014) software, and pericyte number per millimeter (mm) of microvessel length was calculated (24). IgG extravasation was calculated from the 150×150 μm sized ROIs of 8-bit grayscale .tiff format images of wholemount retinas labeled with Cy3-conjugated anti-mouse IgG and lectin. The IgG and lectin frames were applied Huang Thresholding to detect the IgG and lectin-positive pixel counts. After thresholding, the binary images were divided by 255 to convert IgG positive and lectin positive pixels to “1” and background pixels to “0”. Next, the IgG positive pixel counts are divided into lectin positive pixel counts.

### Gene Silencing *in vivo* by Short Interfering RNA (siRNA)

To silence RFC1 gene, we used two custom-designed mouse RFC1 (Slc19a1) Accell siRNAs targeted to two different regions of RFC1 mRNA and two Accell scrambled (control) siRNAs (Horizon Discovery, Waterbeach, United Kingdom). Accell siRNA has several advantages as it does not need any transfection reagents which may have unwanted effects when applied *in vivo*. Furthermore, they could be internalized by any type of mammalian cells (25). Accell siRNAs have been previously proven to be uniformly distributed across rat retinal tissue and successfully inhibit various genes following intravitreal delivery (26). Product codes of siRNAs with sequences are given in Table 1. RFC1-siRNA-1 and RFC1-siRNA-2 or scrambled (control) siRNA-1 and 2 were pooled in equal volumes for administration to improve the potency and specificity.

**Table 1.** The product codes and the sequences of sense strands of RFC1-siRNAs and scrambled (control) siRNAs.

|  | Product codes | Sequences of sense strand |
| --- | --- | --- |
| **RFC1-siRNA-1** | A-044248-13 | C.C.A.G.C.C.U.A.C.U.U.C.A.U.G.C.U.U.U.U.U |
| **RFC1-siRNA-2** | A-044248-15 | C.C.A.G.G.A.A.A.C.U.A.G.A.U.C.G.C.A.U.U.U |
| **Scrambled siRNA-1** | D-001910-01 | U.G.G.U.U.U.A.C.A.U.G.U.C.G.A.C.U.A.A.U.U |
| **Scrambled siRNA-2** | D-001910-03 | U.G.G.U.U.U.A.C.A.U.G.U.U.U.U.C.C.U.A.U.U |

We delivered 100 μM pooled RFC1-siRNAs or Control-siRNAs intravitreally to the eyes (2.3 μL/ eye) of animals via 33 G Neuros Syringe (1701 RN, Hamilton). We did not utilize contralateral eyes as controls to avoid the possible transfer of siRNAs to the opposite side (27, 28). After 24 hr, we either proceeded to retinal ischemia or sacrificed mice, and determined the levels of mRNA or protein.

### Gene Overexpression *in vivo* by Lentiviral Vector (LV)

Lentiviral vectors (LV) that express either green fluorescent protein (GFP) (Control-LV) or GFP with RFC1 (RFC1-LV) driven by EF-1 alpha promoter were used to overexpress RFC1 gene in the retina.

The lentiviral packaging system used for RFC1-LV or control-LV was second-generation and was an effective and safe technique in production. Total RNA from cultured mouse neuro 2a cells was extracted using AllPrep DNA/RNA/Protein Mini Kit (80004, Qiagen, Hilden, Germany) following the manufacturer’s instructions. Transcriptor First Strand cDNA Synthesis Kit (04896866001, Roche, Basel, Switzerland) was used to obtain complementary DNA (cDNA). The coding region of Mus musculus SLC19a1 variant 1/RFC1 (NCBI Reference Sequence: NM_031196.3) was amplified with specific primers (forward 5’-agtcagaattcatggtgcccactggccag-3’and reverse 5’-AGTCAGGATCCTCAAGCCTTGGCTTCGACTCTT −3’) with fast digest restriction enzymes EcoRl (FD0274, Thermo Fisher Scientific, Massachusetts, USA) and BamHI (FD0054, Thermo Fisher Scientific). Restriction enzymes BamHI and EcoRI were used to digest the PCR product of SLC19a1 and expression plasmids (pLenti-EF1α-GFP-2A-Puro; Applied Biological Materials, Richmond, Canada). Subsequently, ligation was realized with T4 DNA ligase (EL0014, Thermo Fisher Scientific). Sequencing pMD2.G and psPAX plasmids, kindly provided by Dr. Didier Trono (Ecole Polytechnique Federale, Lausanne, Switzerland), were used to confirm the insert. They were employed as complementary vectors of the packaging of the lentiviral system (12259; 12260, Addgene, United Kingdom). HEK293T cell line (6 × 10^6^ cells) was chosen to be seeded on 10 cm plates (CLS3294, Corning, New York, USA). The following day, the vector transfection Lipofectamine 3000 (L3000015, Thermo Fisher Scientific) was applied according to the manufacturer’s instructions. Shortly, 7 μg lentiviral vector, 3.5 μg pMD2.G, and 7 μg psPAX were used to prepare DNA-lipid complexes. The DNA-lipid complex was added slowly after 10 min of incubation at room temperature. Six hours after transfection, the medium was changed with fresh DMEM (P04–01158, Pan Biotech, Bavaria Germany) and was incubated at 37°C in a humid atmosphere consisting of 5% CO2. Twenty-four and fifty-two hours following the transfection, the medium was harvested, centrifuged (10 min at 2000 rpm), and filtered with a low binding filter with a 0.45 μm pore size. After ultra-centrifuging (120,000 g for 2 hr), viral particles were dissolved in Dulbecco’s Phosphate Buffered Saline (DPBS) without calcium and magnesium (P04–3650, Pan Biotech). The plasmid with no DNA inserted was packaged with the same procedures to use as a control. The viral titer (1 × 10^8^ lentivirus particles in 1 μl 0.1 M PBS) was measured by a previously published protocol (29).

LV (1 × 10^8^ lentivirus particles in 1 μl 0.1 M PBS) were intravitreally delivered to the retina (2 μL/eye) via 33 G Neuros Syringe (1701 RN, Hamilton). Ten days after the delivery, we either proceeded to retinal ischemia or sacrificed mice to obtain eyeballs.

### Quantitative Reverse Transcriptase-Polymerase Chain Reaction (qRT-PCR)

Eyeballs were freshly dissected in ice-cold sterile PBS under a surgical microscope to get the retina intact. The RNA from whole retinas was extracted via TRIzol Reagent (Invitrogen) according to the manufacturer’s guidelines. The retinal RNA was then reverse transcribed to cDNA with a high-capacity reverse transcription cDNA kit (Applied Biosystems). Primer pairs specific to mouse RFC1/SLC19a1 (Mm00446220_m1) and Tubb4a (Mm00726185_s1, beta 4A class IVA as a housekeeping gene) were constructed and verified by Life Technologies to use with TaqMan qPCR chemistry (TaqMan™ Gene Expression Assay, Applied Biosystems™, 4331182). Amplicon context sequences for SLC19a1 and beta 4A class IVA are shown in Table 2. qRT-PCR analysis was performed to calculate the relative fold change in mRNA expression level of the SLC19a1 after siRNA administration. The critical threshold cycle (CT) of SLC19a1 gene was first normalized to the beta-tubulin 4A with the comparative CT method. The ΔCT which is the difference in CT values between the SLC19a1 and beta-tubulin 4A genes were then normalized to the corresponding ΔCT of the control siRNA group, expressed as ΔΔCT and calculated as fold change (2^−ΔΔCT^).

**Table 2.**
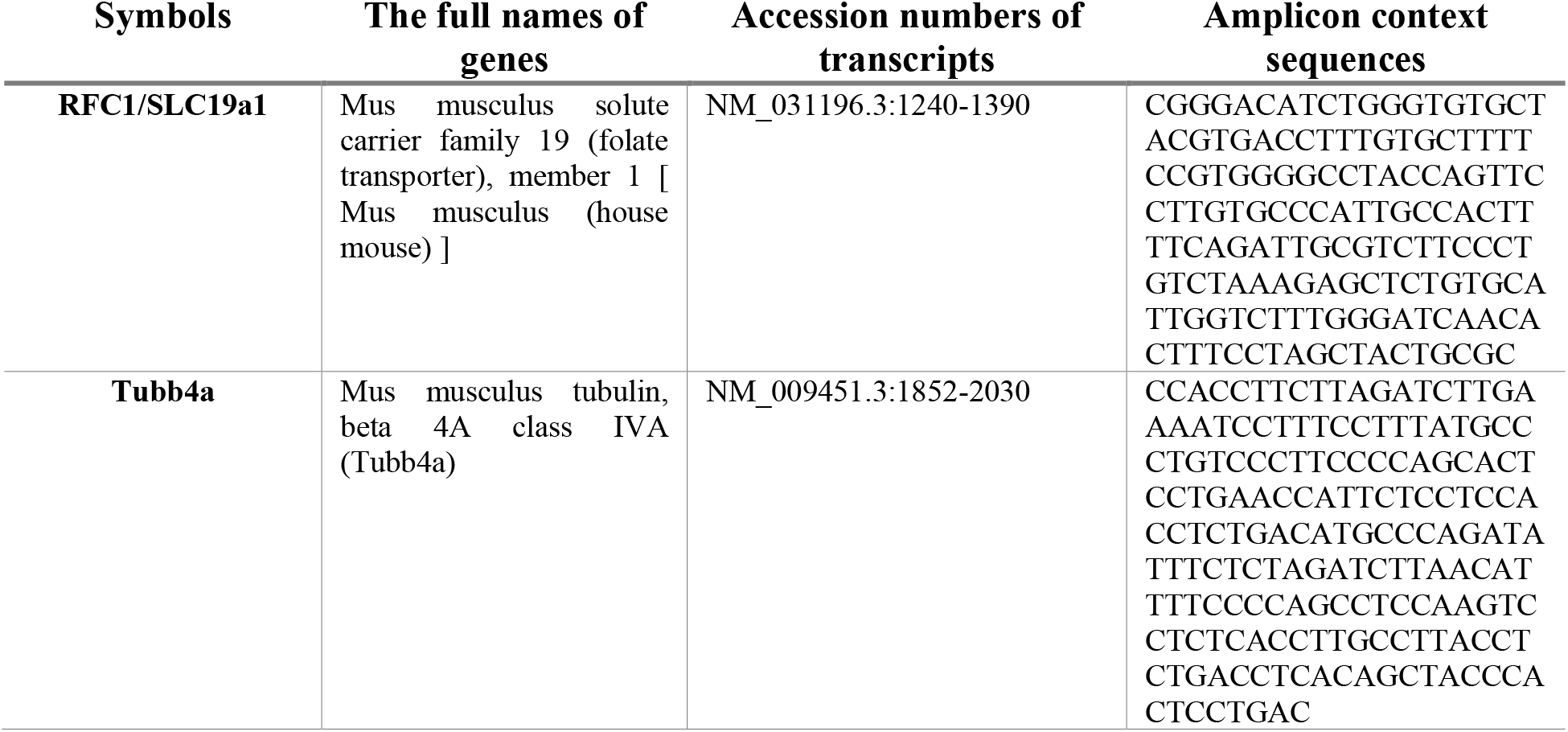
Gene symbols, the full names of the genes, accession number of transcripts and amplicon context sequences of SLC19a1 and beta 4A class IVA.

### Retinal Ischemia Model

The retinal ischemia model used here was previously developed in our laboratory (30). In brief, anesthetized mice were put in a prone position under a stereoscope (SMZ1000, Nikon Instruments Inc., Amsterdam, The Netherlands) and their heads were immobilized with a nosepiece. Central retinal arteries located in the central optic nerve were exposed via cautious retroorbital dissection. A small strip (0.3 × 1 mm) of 30% FeCl_3_–soaked filter paper was placed on the optic nerve for 3 min to trigger clot formation and occlude the central retinal artery. Based on our previous observations, we used 1 hr of ischemia, as it was sufficient to induce ischemic changes in the microcirculation such as mural cell contractions (18, 19). One hr later, eyeballs were collected under anesthesia, and animals were sacrificed by cervical dislocation.

### Statistical Analyses

All in vivo experiments were repeated in at least n=3 mice/group. The number of mice per group for *in vivo* experiments are mentioned in the figure legends. All results were conveyed as mean ± standard error of the mean (S.E.M.). Data were analyzed using IBM SPSS 23. Non-normally distributed data were compared using the Mann-Whitney U test (for two groups). The expression ratio in a treatment group was expressed as a percentage of the ratio in the control group. p≤0.05 was considered significant.

## RESULTS

### RFC1 protein is abundantly expressed in the endothelial cells and pericytes of the inner BRB

The scarcity of antibodies for RFC1 immunohistochemistry (21) until recently was a challenge for studies; hence, most former studies used RFC1 antiserum or homemade antibodies to mark RFC1 protein in the tissues (21, 31, 32). The availability of commercial RFC1 antibodies that were confirmed by immunohistochemistry and by Western Blotting (1, 2) facilitated the research in the field.

We first immunohistochemically examined RFC1 in the coronal retina sections, and as expected, determined that RPE was immunopositive (Fig. 1A). This was also confirmed in whole mount retina preparations (Fig. 1B) (33). RFC1 immunopositivity was also found in the ganglion cell layer (GCL), inner and outer plexiform layers (INL and OPL) that coincided with the superficial and deep microvascular plexus. Hence, the RFC1 immunosignal was identified in the retinal layers comprising microvessels (Fig. 1A, C). We used two different antibodies, high concentrations, and long incubation durations like three overnights at +4°C with the primary antibody for the 200 μm thick whole mount retinal preparations to visualize retinal microvessels. Likewise, isolating retinal microvessels with the trypsin digestion method provided us with an easier and more consistent preparation for immunostainings.

**Fig. 1.**
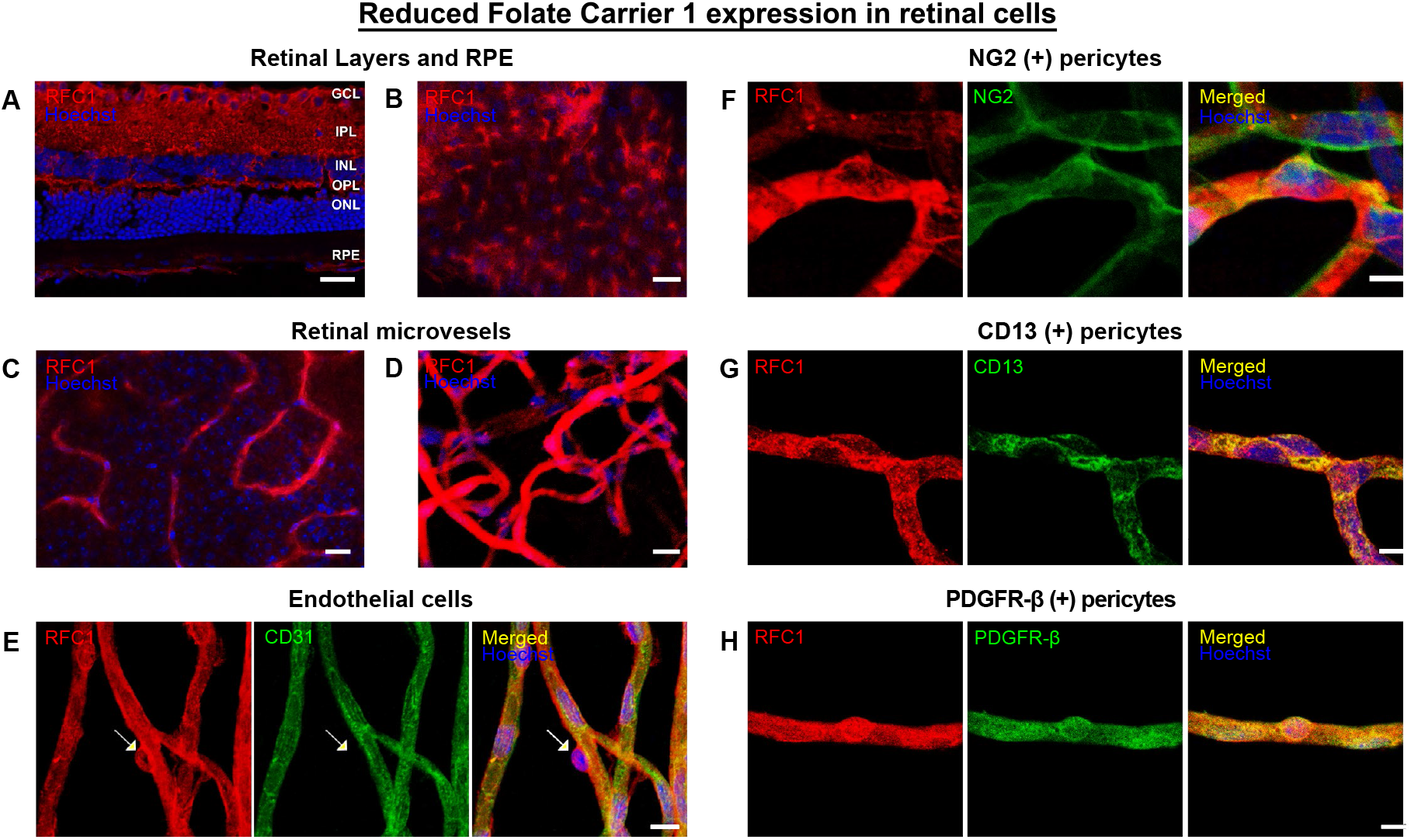
RFC1 protein is abundantly expressed in endothelial cells and pericytes of the retinal microvessels. A) Ex-vivo labeling of 20 μm coronal cryo-section from PFA fixated eyeballs of naive Swiss Albino mice with anti-RFC1 antibody (red). Abundant RFC1 immunopositivity is observed along ganglion cell layer (GCL), inner plexiform layer (IPL), outer plexiform layer (OPL) where retinal microvessels forms vascular horizontal vascular beds; as well as the retinal pigment epithelium (RPE). However, this preparation limited the observation of microvessels, hence inner BRB, as it commonly included microvessels circularly rather than longitudinally. Scale Bar=25 μm B) RPE which is known to express RFC1 previously is well stained with anti-RFC1 antibody (red) as our positive control. C) The microvessels constituting inner BRB form the deep vascular plexus of the retina were immunohistochemically labelled with anti-RFC1 antibody in PFA fixated whole-mount retinas D) Retinal microvessels which were obtained via retinal trypsin digestion method that allowed to get only microvessels (< 9 μm diameter) were immunofluorescently labeled with anti-RFC1 antibody (n=6 retina; red). Nuclei were labeled with Hoechst 33258 (blue) in all the rows. E) the endothelial marker CD31 (green) (n=3). White arrow show “bump-on-a log” shaped pericyte body was positively stained with anti-RFC1 antibody, but negative for endothelial marker CD31. Hence, RFC1 staining was specific. F-H) RFC1 (red) also colocalized with accepted pericyte markers NG2, CD13, PDGFR-β (green) shown respectively (n=3/marker). Nuclei were labeled with Hoechst 33258 (blue). Scale bar: 10 μm.

We observed that RFC1 was distributed continuously and diffusely throughout the retinal microvessels (Fig. 1D). RFC1 was immunohistochemically co-labeled with the endothelial marker, cluster of differentiation 31 (CD31), whose immunoreactivity exclusively outlined the endothelial cells. As a convincing finding that there was no crosstalk in immunoreactivity, RFC1 labeling was determined to colocalize with both the endothelial cells, and the ‘bump on a log’ shaped cells (Fig. 1E, arrows), which designates pericytes.

We then labeled the microvessels with well-defined pericyte markers including the neural/glial antigen 2 (NG2), aminopeptidase-N (CD13), and platelet-derived growth factor receptor beta (PDGFR-β) all of which showed colocalization with RFC1 (Fig. 1F-H). These indicated that RFC1 protein was present both in the endothelial cells and pericytes.

### Retinal RFC1 is downregulated *in vivo* by RFC1 targeted Accell siRNA, and leads to disruption of the inner BRB

We delivered a combination containing the same amount of specifically designed two RFC1 targeted Accell siRNAs (RFC1-siRNA) or two scrambled siRNAs intravitreally and sacrificed the mice 24 hr later. To confirm the knockdown of RFC1, we obtained fresh whole retinas and determined a pronounced decrease in RFC1 protein via Western Blotting (p= 0.028; Fig. 2A, B). Additionally, RFC1-siRNA significantly decreased RFC1 mRNA by 24.75% compared to scrambled siRNA (p= 0.004, Fig. 2C). In line with these, RFC1 immunosignal intensity was diminished in the microvessels of RFC1-siRNA injected retinas when mean grey value of RFC1 immunosignal (i.e., fluorescence intensity) was measured (72.1%, p<0.0001; Fig. 2C, D). We also noticed that RFC1-siRNA treated retinas were fragile, and prone to damage during the procedures.

**Fig. 2.**
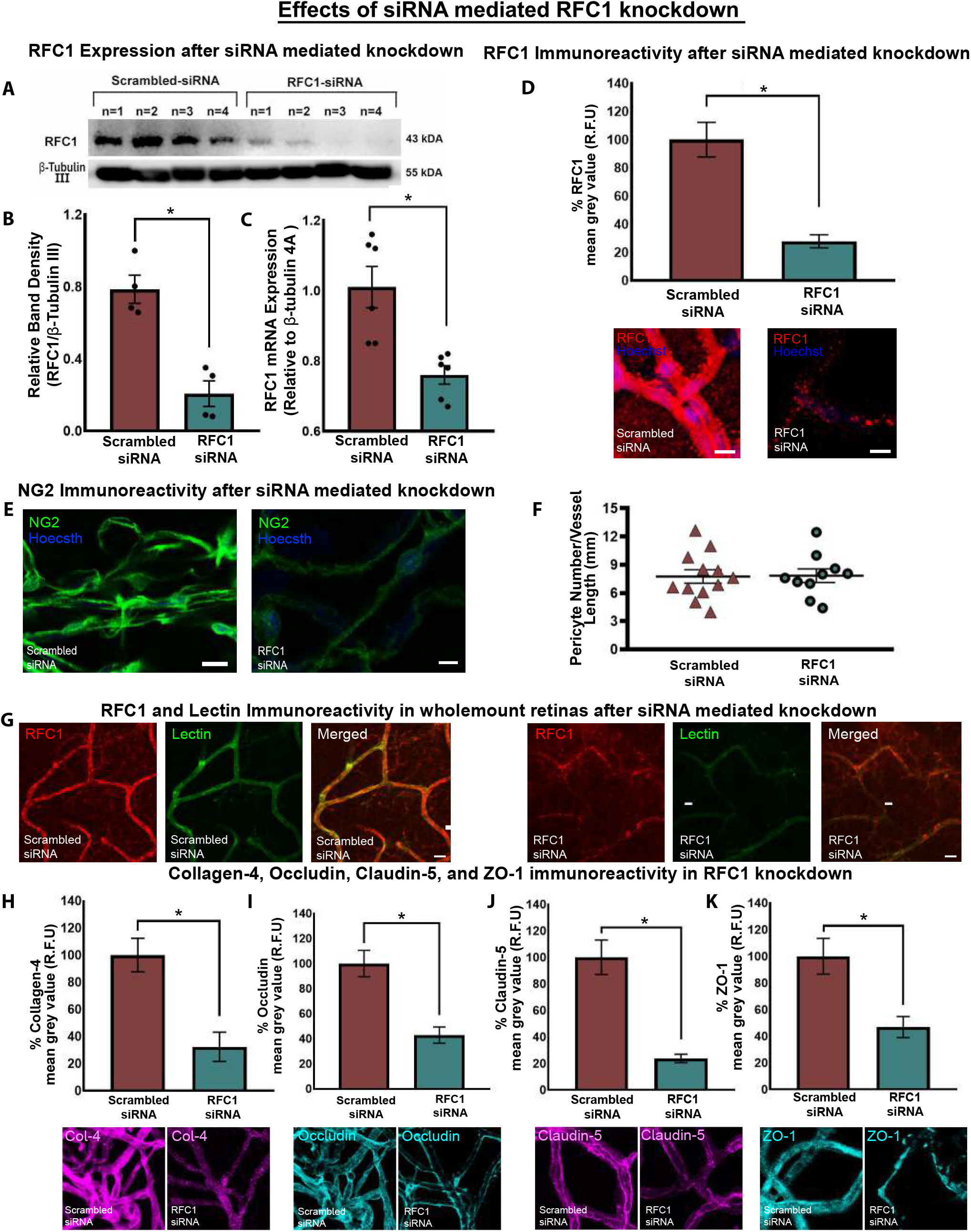
The validation of Accell siRNA mediated *in vivo* RFC1 knockdown, which led to a reduction in critical proteins of the inner BRB. A) Western blotting image of retinas (n=4 mice/group) depicting the robust decrease in RFC1 protein of RFC1-siRNA delivered ones compared to scrambled-siRNA. β-Tubulin III was loading control. 35 μg protein was loaded for each retina. B) The graph illustrates the quantification of the relative band densities of given Western blotting image which expresses RFC1 band density in proportion to β-Tubulin III, which shows decrease in relative protein levels in RFC1-siRNA treated retinas (*p= 0.028). C) The graph shows that RFC1-siRNA delivery reduced retinal RFC1 mRNA levels by 24.75% when compared to scrambled-siRNA (*p=0.004). D) The graph illustrates the percentage of mean grey value of RFC1 frame in lectin positive microvessel area in RFC1-siRNA treated group normalized to Scrambled-siRNA treated group as described in Methods section. RFC1-siRNA delivery significantly reduced the percent of the mean grey values of RFC1 by 72.10% (n=4). Representative confocal images of Scrambled-siRNA or RFC1-siRNA delivered retinal microvessels stained by anti-RFC1 antibody (red). E) RFC1-siRNA treated retinal microvessels showed less NG2 immunosignal compared to Scrambled-siRNA treated ones, indicating that pericytes might also be damaged by RFC1-siRNA. F) However, pericyte body counts per microvessel length (mm) were not different between the groups. n = 3 images were analyzed per animal. G) Whole mount retinas were labeled by anti-RFC1 antibody (red) and Fluorescein Lectin (green). RFC1 immunosignal was decreased and interrupted along deep retinal microvessels. Also, lectin signal was weakened in RFC1-siRNA treated retinas representing the structural decomposition of microvessels. H-K) RFC1-siRNA delivery significantly reduced the percent of the mean grey values of Collagen-4 68% (n=4), Occludin 57.03% (n=3), Claudin-5 76.26% (n=4), and ZO-1 53.22% (n=4). The reductions that are statistically significant represented as *p ≤ 0.05. Immunosignal decrease was also observed collagen-4 (magenta), Occludin (cyan), Lectin (Yellow) as well as G, H) Claudin-5 (magenta), and ZO-1 (cyan) via RFC1-siRNA treatment compared to Scrambled-siRNA treatment. Nuclei are labelled with Hoechst 33258 (blue), Data are mean ± S.E.M Mann-Whitney U; Scale bar: 10 μm.

The immunostaining of the whole mount retinas also confirmed RFC1-siRNA knockdown with diminished RFC1 immunosignal, and further showed that Lectin staining was weaker and interrupted in RFC1-siRNA treated preparations, indicating that microvessels might be structurally damaged (Fig. 2G). In addition, these retinal microvessels demonstrated lower NG2 signal compared to scrambled siRNA treated ones, which suggested that pericytes might also be damaged by RFC1-siRNA (Fig. 2E). However, to address if the reduction in RFC1 immunoreactivity in siRNA treated microvessels might be associated with the loss of pericytes, we assessed pericyte density (soma per millimeter capillary). We found no significant decline in pericyte density in RFC1-siRNA administered retinas suggesting that RFC1 decrease may not affect pericyte maintenance, as it did not reduce pericytes in number at least at this time point (Fig. 2F). However, we could not exclude disruption of pericyte functions via RFC1-siRNA since there is marked decline in the expression of pericyte marker NG2.

As we determined that retinal microvessels express RFC1 protein abundantly, and we could efficiently knock down RFC1 *in vivo*, we elucidated whether RFC1 suppression may lead to any changes in the inner BRB.

Strikingly, knocking down retinal RFC1 diminished immunoreactivity of the tight junction-associated transmembrane proteins occludin, claudin-5, and cytoplasmic adaptor protein zonula occludens-1 (ZO-1), as well as the main basement membrane protein collagen-4 in trypsin digested microvessels (Fig. 2H-K). In RFC1-siRNA treated microvessels, immunoreactivity of collagen-4 that surrounds the abluminal membrane of endothelial cells and covers pericytes residing in the basement membrane decreased by 68% (p=0.0043), occludin decreased by 57.03% (p=0.0043), and claudin-5 decreased by 76.26% (*p*= 0.0006). ZO-1, the intracellular adaptor protein establishing a link between occludin and intracellular actin cytoskeleton decreased by 53.22% (p= 0.0012).

### Retinal RFC1 is upregulated *in vivo* by LV overexpressing RFC1, and this upregulation changes the inner BRB properties

Next, we aimed to increase the expression of RFC1 in mouse retinas via LV carrying RFC1 as described in detail in the methods section. We preferred the LV gene delivery method due to its efficiency to induce stable gene expression, its tropism to the inner retina and the cells of retinal microvessels when delivered intravitreally (34, 35).

The efficient RFC1 protein overexpression was shown by Western Blotting in cell lysates (Fig. 3A). Subsequently, we proceeded to validate the *in vivo* LV-mediated transduction. Mice had unilateral intravitreal LV injections. No animals developed any local infections. The eyeballs of mice (n=3) that received intravitreal Control-LV bearing GFP were harvested 10 days after. The retinal and perimicrovascular cells showed GFP expression disclosing that the cells were infected with LV (Fig. 3B).

**Fig. 3.**
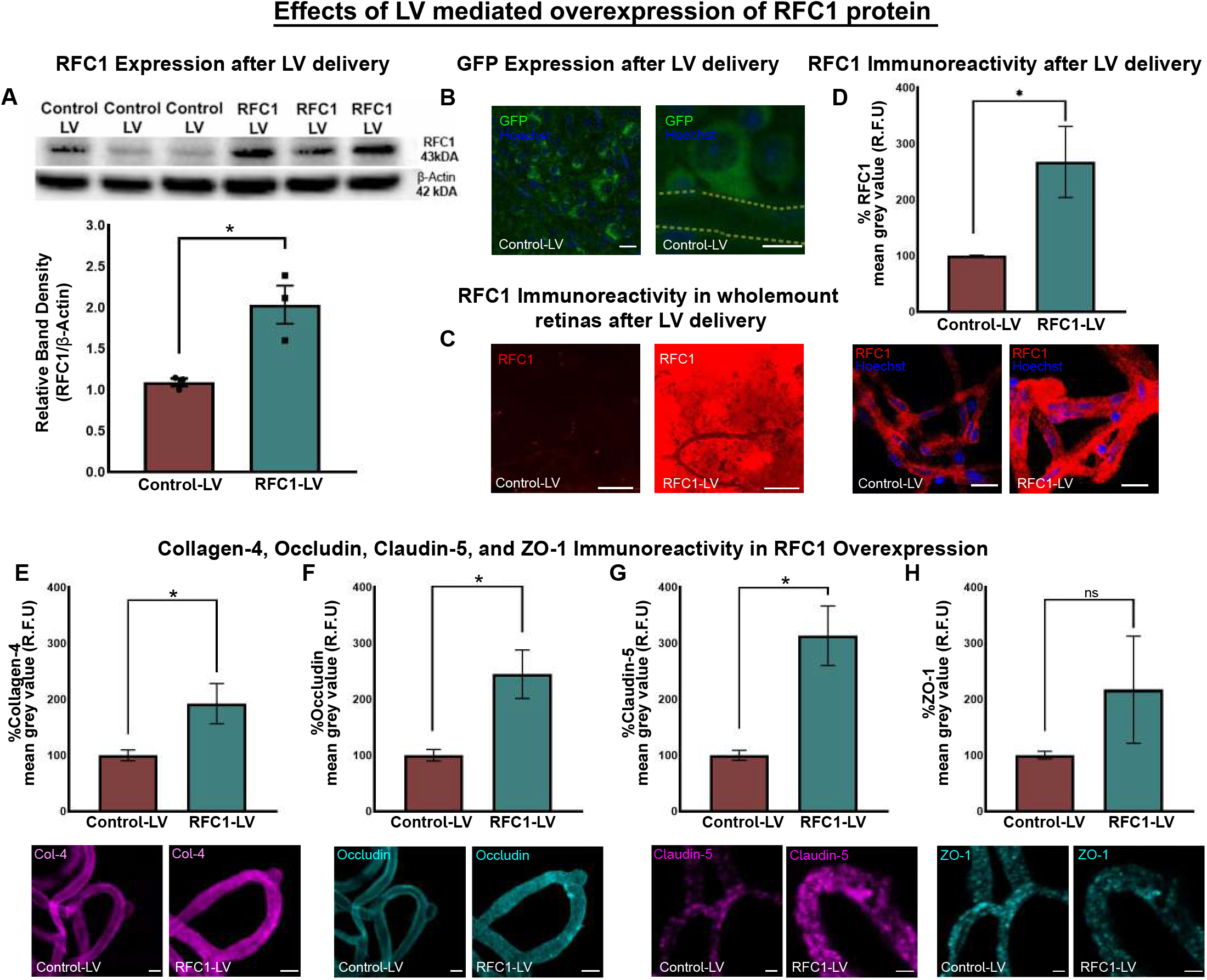
The validation of lentiviral vector-mediated overexpression of RFC1 protein *in vivo*, which led to an increase in the proteins of BRB. A) The Western Blotting image shows corresponding overexpression of RFC1 protein in RFC1-LV transduced neuroblastoma cells (N2a) compared to Control-LV transduced cells. For cell culture lysates, Beta-Actin was the loading control. B) The graph illustrates the quantification of the relative band densities of given Western blotting image which expresses RFC1 band density in proportion to β-Actin, which shows increase in relative protein levels in RFC1-LV treated retinas (*p= 0.016). B) The confocal images from whole mount retinas confirmed the intravitreal delivery of Control-LV into the mouse eye showing lentiviral vectors infected various retinal cells, as reporter gene GFP (green) signal indicated. Also, as magnified image showed, perivascular cells and pericytes expressed GFP protein in LV-RFC1 or Control-LV injected retinas, microvessel trace was shown by dashed line. C) The representative images of whole mount retinas which had been treated by LV-RFC1 showed increased RFC1 immunosignal (red) compared to Control-LV injected control retinas (n=3). D) The graph illustrates the percentage of mean grey value of RFC1 frame in lectin positive microvessel area in RFC1-LV treated group normalized to Control-LV treated group as described in Methods section. RFC1-LV delivery significantly increased the percent of the mean grey values of RFC1 (n=3). E-H) RFC1-LV delivery significantly increased the percent of the mean grey values of Collagen-4 (n=4), Occludin (n=3), Claudin-5 (n=3), but not ZO-1 compared to Control-LV delivered groups. (*p ≤ 0.05). Collagen-4 (magenta), Occludin (cyan) Lectin (Yellow) immunosignal increased as well as Claudin-5 (magenta) except ZO-1 (cyan) via RFC1-LV treatment compared to Control-LV treatment. Nuclei were labelled with Hoechst 33258 (blue) Data are mean ± S.E.M. Mann-Whitney U; Scale bar: 10 μm.

The overexpression of RFC1 by LV in the retina was also observed immunohistochemically. RFC1 immunoreactivity revealed a marked intensity increase in both the retinal whole mount and retinal microvessel preparations in RFC1-LV compared to Control-LV (Fig. 3C, D) with 167.5% increase in RFC1 immunosignal (p= 0.0159; Fig. 3D).

As the suppression of RFC1 levels led to an extensive structural damage in retinal microvessels, we further investigated whether RFC1 overexpression had any effects. Immunohistochemistry of tight junction proteins (occludin, claudin-5), intracellular adaptor protein ZO-1, and main basement membrane protein collagen-4 in retinal microvessels (from n=3 retina/per marker, Fig. 3E-H) were done ten days after administration. RFC1-LV administered retinal microvessels displayed increased immunosignal in occludin (p= 0.0059), claudin-5 (p= 0.0040), and collagen-4 (p= 0.0159) compared to Control-LV ones (Fig. 3E-G). However, the immunosignal of ZO-1 did not differ between the groups (Fig. 3H).

### Retinal ischemia alters RFC1 protein and decreases the inner BRB proteins which can be ameliorated by RFC1 overexpression

After observing the potential role of RFC1 protein in maintaining the inner BRB, we investigated whether RFC1 protein has a role in retinal ischemia. We made 1 hr permanent retinal ischemia, as it is considered sufficient to observe microvessel-related changes such as capillary constrictions or protrusion of the contracted pericyte soma from the microvessel wall (Fig. 4C, white asterisk), and yet an early time point to allow us to minimize the effects of inflammation (18–20).

**Fig. 4.**
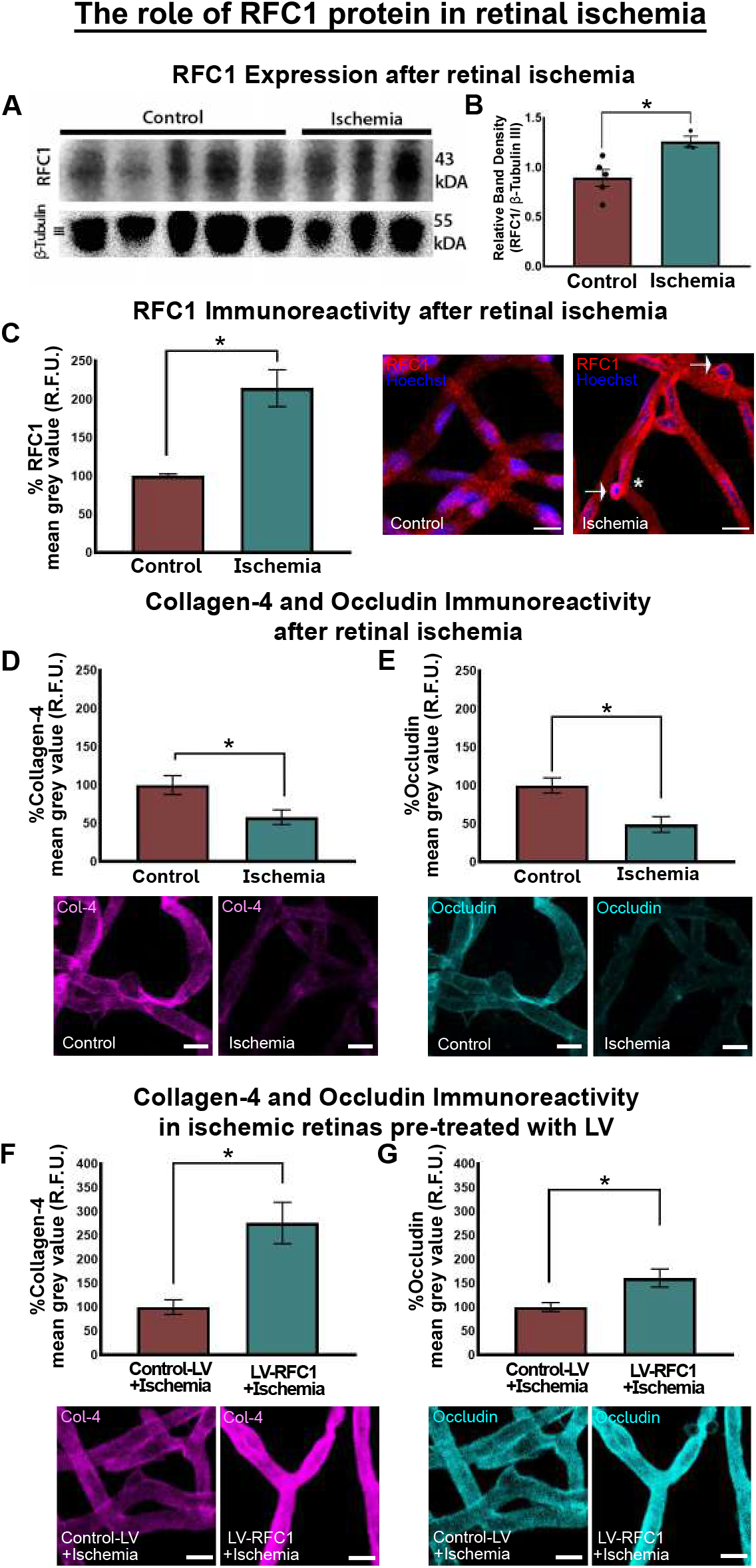
RFC1 protein increased after retinal ischemia, and the overexpression of RFC1 before ischemia by LV salvaged decreased collagen-4 and occludin levels. A) The representative Western blotting image from Control (n=5) and ischemic (n=3) retinas. Housekeeping gene β-Tubulin III was used as loading control. B) The graph shows the relative density measurements of bands which were calculated by proportioning to loading control. One-hour permanent retinal ischemia increased relative RFC1 protein levels compared to control. C) Retinal ischemia significantly augmented the percentage of mean grey value of RFC1 by 114.5% (n=3). Representative confocal images of control or ischemic retinal microvessels stained by anti-RFC1 antibody (red). Significant immunosignal increase in ischemic microvessels was observed compared to controls. Also, ischemic microvessels demonstrated expected characteristics such as constrictions and protruding pericyte bodies (asterisk). Of note, ischemic pericyte bodies showed denser RFC1 immunosignal (white arrows). D, E) In contrast, retinal ischemia decreased collagen-4 by 42.07% (n=3), Occludin by 50.94% (n=3). F, G) In addition, microvessels treated with LV-RFC1, 10 days before ischemia showed 176% increase in Collagen-4 (n=3) and 60.9% in Occludin (n=3) immunosignal compared to Control-LV delivered ones indicating RFC1 overexpression before ischemia might retrieve loss of collagen-4 and occludin in ischemia. (*p ≤ 0.05). Nuclei were labelled with Hoechst 33258 (blue) in all images. Data are mean ± S.E.M. Mann-Whitney U; Scale bar: 10 μm.

Following ischemia, the expression of RFC1 protein was enhanced significantly compared to controls detected by Western blotting (p= 0.025; Fig. 4A, B) and RFC1 immunopositivity (p <0.0001; Fig. 4C). The pronounced RFC1 signal was detected in the protruding pericytes (Fig. 4C, white arrows) probable of an increased signal in affected cells.

We studied whether retinal ischemia caused any alterations in the inner BRB proteins and observed that 1 hr retinal ischemia decreased collagen-4 and occludin immunosignal compared to controls (*p* =0.0140, *p* =0.0317 respectively; Fig. 4D, E).

Finally, we studied whether RFC1-siRNA knockdown induced disruption of the inner BRB proteins resulted in barrier dysfunction. We stained vessels with lectin and determined endogenous IgG extravasation by fluorescently labeled anti-mouse IgG antibody (Fig. 5A, red arrows). RFC1-siRNA treated mice displayed endogenous IgG leakage (Fig. 5A, red arrows), while scrambled-siRNA treated retinas did not show any. This indicated that RFC1 is essentially required for the structural and functional integrity of the inner BRB in health.

**Fig. 5.**
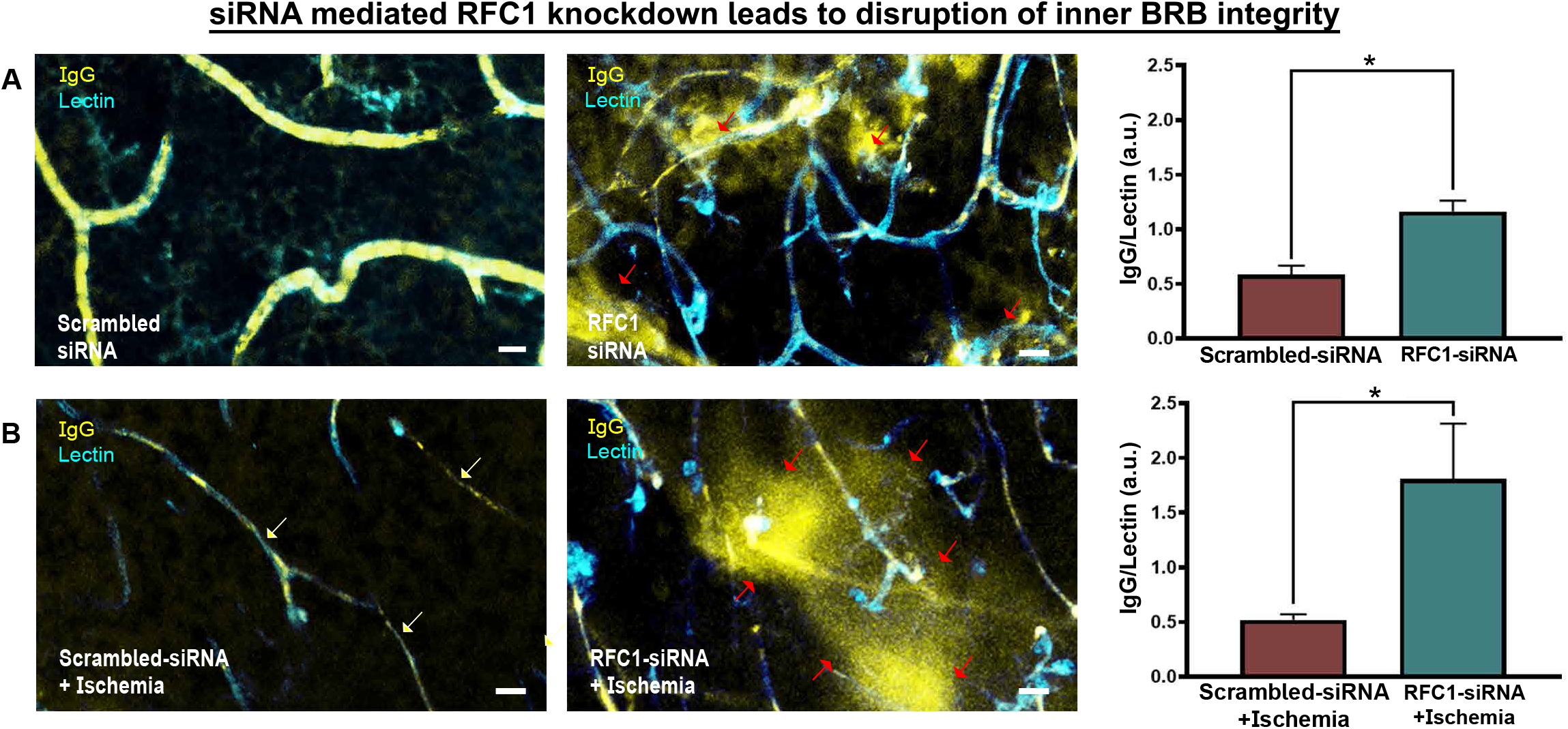
siRNA mediated RFC1 knockdown led to inner BRB breakdown and endogenous IgG extravasation. The wholemount retinas were labeled by Lectin (cyan) and incubated with Cy3 labeled goat anti-mouse antibody (yellow) to visualize the extravasated endogenous mouse IgG. A) RFC1-siRNA mediated knockdown led to significant IgG extravasation (red arrows) in microvessels while B) Control-siRNA treated retinas showed no extravascular IgG signal (p=0.001; n=3). C) 1 hr ischemia led to microvessel constrictions (white arrows) but no IgG extravasation. D) However, knocking down RFC1 before ischemia provoked inner BRB disruption leading to endogenous IgG extravasation (red arrows) (p=0.05; n=3). Nuclei were labelled with Hoechst 33258 (blue) in all images. Scale bar: 10 μm

On the other hand, we observed no IgG extravasation after 1 hr retinal ischemia, so it significantly did not compromise the integrity of inner BRB (Fig. 5B). This observation was consistent with the literature since 1 hr retinal ischemia causes early changes such as capillary constrictions (white arrows, Fig. 5B), but it does not suffice for the inner BRB disruption to the extent of allowing endogenous IgG (150 kDa) leakage through paracellular barrier even if disintegration of tight junctions commenced (36–38). We then studied whether knocking-down RFC1 before ischemia may provoke disruption of the inner BRB. When RFC1-siRNA was administered 24 hr before retinal ischemia, a significant amount of endogenous IgG extravasation was determined (Fig. 5D, red arrows). As scrambled siRNA administration 24 hr before retinal ischemia led to no apparent IgG extravasation, this indicated that decreased RFC1 levels accelerated the inner BRB disruption in acute retinal ischemia.

## DISCUSSION

In this report, for the first time, we demonstrated the presence of RFC1 protein in the endothelial cells and pericytes of the inner BRB. We also showed that RFC1 - once considered only as a transporter protein - is an essential component for maintaining the inner BRB and has fundamental roles in pathological conditions affecting the inner BRB, such as retinal ischemia.

RFC1 expression at the protein level in endothelial cells and pericytes of retina has never been studied before, despite its consistent detection in transcripts of microvascular mural cells (pericytes and vascular smooth muscle cells) of the brain (39). Recently, a study characterized the localization of folate transporters, hence RFC1 expression at CNS barriers in detail focusing especially on BBB cells including the endothelial cells; however, this study explored neither the retina nor pericytes (40). So, our study bears another focus, and offers a recognition for the roles of RFC1 in the CNS and deepens our understanding of other critical functions as well as novel treatment strategies for brain disorders.

Isolating retinal microvessels with the trypsin digestion method provided us an easier and more consistent preparation for immunostainings. Our results highlight that, retinal endothelial cells, like their counterparts in the CNS, express RFC1 protein. As an additional novel finding we detected the colocalization of RFC1 with the well-defined pericyte markers. Most studies evaluating the role of RFC1 in microvessels concentrated on only the endothelial cells, but not the neighboring pericytes (1, 2, 31, 41). In this regard, RFC1 protein was never characterized in CNS pericytes as well. Our findings actually match the results of a recent meta-analysis combining five different mouse brain pericyte transcriptomic studies where the RFC1 gene was one of the only three consistently detected genes in brain pericytes out of the 1180 enriched genes (39).

Our study provides some initial insights regarding the functional importance of RFC1 in retinal microvessels. The genetic knockout of RFC1 is lethal (42) and no conditional knockout animals had been used in the studies before, so we decided to utilize siRNA technology to manipulate RFC1 levels specifically in the retina of adult mice. Two previous *in vitro* studies using RFC1-siRNA which were performed in primary cultures of human differentiated adipocytes (43), and in rat choroidal epithelial Z310 cells (44), and both did not focus on the microvessels or the retina. Hence, our method to knockdown RFC1 *in vivo* by RFC1 targeted custom designed Accell RFC1-siRNA could be considered as one of the initial studies. RFC1-siRNA administration to the retina *in vivo* via intravitreal delivery offers some advantages: 1) the retina is the easily reachable, well-known extension of the brain 2) the vitreous cavity is an enclosed space that minimizes the adverse effects of systemic spread and maximizes the effect of applied concentration 3) delivery is fast and feasible (45). The application of RFC1-siRNA deranged the structure of retinal microvessels, specifically the deep plexus comprising blood barrier properties, as shown by lectin stainings (Fig. 2E). Interestingly, knocking down RFC1 led to a significant decrease in the barrier proteins occludin, claudin-5, intracellular adaptor protein ZO-1, and the main basement membrane protein collagen-4. This induced the inner BRB disintegration and led to functional failure of the barrier properties. Although it may be plausible to interpret this striking result with caution, there is a study showing that the tight junction proteins like occludin, claudin-1, and ZO-1 were substantially decreased in the BBB of capillaries isolated from proton coupled folate transporter (PCFT) null mice in comparison to controls (46). However, this mouse with deleted PCFT gene - another functionally distinct folate transporter- is accepted as a model for hereditary folate malabsorption that leads to systemic folate deficiency. Hence, it is not possible to fully exclude the contribution of chronic folate deficiency to these effects. However, in our study, we observed inner BRB changes after acute modifications of the transporter in naive mice, which circumvents the confounding developmental effects of folate deficiency. In line with these observations, our experiments with the retinal overexpression of RFC1 via LV have shown an upregulation of occludin, claudin-5 and collagen-4, supporting the role of RFC1 in inner BRB protein expressions. As an interesting observation, RFC1 was recently determined as a cGAMP importer in the cyclic GMP–AMP synthase (cGAS)–stimulator of interferon genes (STING) pathway that has a role in inflammation, infection, cellular stress, tissue damage, and tumor angiogenesis (21, 47, 48). In mouse tumor models, intratumoral cGAMP treatment led to a 40% decrease in the density of the vessels, a 1.7-fold rise in pericyte coverage, and a 1.5-fold rise in collagen-4 coverage (47), which may indicate that RFC1 may indirectly involved in remodeling the tumor vasculature in consistent with our observations in inner retinal microvasculature.

Based on reports that show pericytes are eminent for BBB and inner BRB maintenance and contribute to tight junction formation and preservation (49–52), we also excluded pericyte deficiency in RFC1-siRNA treated retinas. Although we found no significant decrease in pericyte density by counting pericyte soma per mm capillary in RFC1-siRNA treated mice, we observed immunosignal decrease in pericyte marker NG2, which involves in many functions of pericytes including proliferation, motility and importantly, endothelial cell junction assembly by activating integrin signaling (53). For example, the treatment of human microvascular pericytes with NG2-siRNA hindered collagen-4 coverage, ZO-1 expression, and endothelial junction maintenance in endothelial cells. Likewise, tumors from mice whose pericytes do not express NG2 showed similar changes in their endothelial junctions in the same study (28). Although the underlying mechanisms require clarification, we suggest that NG2 might be a mediator of the effects observed in tight junction proteins and collagen-4 via RFC1-siRNA. Alternatively, the disintegration of NG2 in RFC1-siRNA treated retinal microvessels may indicate more comprehensive dysfunction throughout the inner BRB, to which NG2 disruption is only a contributing factor.

Aside from its role in the maintenance of inner BRB under physiologic conditions, we further investigated the effect of RFC1 in acute retinal ischemia. Retinal ischemia which is a major underlying condition of blindness worldwide is a severe condition with a poor prognosis generally caused by acute occlusion of retinal arteries (54). We induced retinal ischemia by an established method in our laboratory (19, 30), and showed an increase in RFC1 protein after 1 hr permanent ischemia. A previous gene profiling study showed RFC1 mRNA upregulation following 1 hr ischemia / 24 hr recanalization induced by high intraocular pressure retinal ischemia model in rats (55). In contrast, the same study observed no significant change in RFC1 mRNA in 1 hr permanent retinal ischemia. Another study showed that 45 min retinal ischemia followed by 48 hr recanalization upregulated RFC1 mRNA in rats (56). The possible explanation for these results may be that the retinal ischemia models were different, and the authors only determined mRNA levels, not the protein levels of the samples. The estimation of protein levels from mRNA levels might be unreliable, as post-translational changes could affect the protein levels in early time-points (57). There is only one study performed by kidney ischemia-reperfusion model that determined RFC1 protein levels decreased *in vivo* focusing on the proximal tubules where folate reabsorption occurs, emphasizing its importance in adjusting serum folate levels (58). Contrarily, the increase of metabolic needs due to acute phase of ischemia might necessitate the upregulation of RFC1 after 1 hr of retinal ischemia for the preservation of the barrier properties. In line with that, further augmentation of RFC1 expression by LV-RFC1 intervention before ischemia salvaged the decreased occludin and collagen-4 levels, otherwise which would be attenuated by ischemia.

Also, knocking down RFC1 prior to ischemia impaired barrier functions which were demonstrated by endogenous IgG leakage, supporting our findings that RFC1 is essential for the protection of inner BRB during retinal ischemia. This result might be interpreted in the light of the human meta-analysis showing that the SNPs of RFC1 gene differed between ischemic stroke and control groups, and some genotypes were found to be associated with small vessel occlusions and silent brain infarctions (59). Since the changes in RFC1 function or structure with these polymorphisms had not been defined, it is possible that our finding of changes in inner BRB via the genetic modifications of RFC1 could carry the potential of being the experimental evidence of the importance of RFC1 in ischemia and small vessel disease. Thus, intriguing questions regarding the therapeutic potential of intervening RFC1 before, during or after ischemia rises.

Our study can be considered as an initial step to investigate the role of ‘once an unnoticed transporter protein’ in the inner BRB. We took a step-by-step approach, defined the presence of the RFC1 protein in the retinal microvessels and established *in vivo* genetic tools to clarify the potential new roles of RFC1 under physiologic conditions, ischemia. However, some limitations of our study merit consideration. The genetic manipulations we used were not targeted to cells. In addition, we did not study the underlying pathways of how RFC1 regulates barrier proteins or BRB integrity, and the interplay of folate and other folate transport systems (FRα and PCFT) or the concert of the efflux systems (Pgp/ABCB1, MRPs/ABCC, BCRP/ABCG2) (60), which warrants further investigations.

Despite these limitations, our data suggest several implications. We propose a conceptual model of workflow for the other proteins like RFC1 whose presence and roles have not been elucidated so far. Nowadays, the roles of these proteins have been investigated through advanced mathematical models, pathway analyses and the -omics approaches, but the old phrase “seeing is believing” should not be underestimated and *in vivo* studies may provide surprising insights into the role of a protein. The upregulation of RFC1 may be attempted for diseases other than ischemia where the impairment of BRB or BBB is involved in the pathophysiology. Although merely an initial step, our results are encouraging as the presence of RFC1 in inner BRB may be exploited for targeted drug delivery such as RFC1-targeted nanodrugs, folate conjugated nanodrugs or even clinically widely used RFC1 substrate, methotrexate, can be exploited not only in ischemia, but in diseases like cancer.

## CONCLUSION

In the present study, we investigated the presence of RFC1 in the retinal endothelial cells and pericytes, and its role in the regulation of inner BRB integrity. For the first time, we introduced and discussed the tools for the manipulation of RFC1 levels or function *in vivo* which included RFC1-targeted Accell siRNA to knockdown RFC1, and LV to overexpress RFC1. By using these tools, we tried to elucidate the role of RFC1 in the abrupt regulation of inner BRB during physiological conditions and acute retinal ischemia and highlighted its potential role for preserving BRB integrity. This study aimed to precede the further investigation of RFC1 in other tissues and diseases by suggesting a conceptual and methodological framework. Future research may extend this work by integrating pathways which RFC1 is associated with to address the concern of how RFC1 regulates inner BRB.

## Abbreviations

BBB: blood-brain barrier
BRB: blood-retina barrier
CD13: Aminopeptidase N
cGAMP: Cyclic guanosine monophosphate–adenosine
cGAS: Cyclic GMP-AMP synthase
GCL: ganglion cell layer
IgG: Immunoglobulin G
INL: inner nuclear layer
IPL: inner plexiform layer
LSCI: Laser speckle contrast imaging
LV: Lentiviral vectors
mRNA: messenger RNA
MTX: Methotrexate
NG2: Neural/glial antigen 2
ONL: outer nuclear layer
OPL: outer plexiform layer
PBS: phosphate buffered saline
PDGFR-β: platelet-derived growth factor receptor beta
PFA: paraformaldehyde
qRT-PCR: Quantitative Reverse Transcriptase-Polymerase Chain Reaction
RBF: Relative blood flow
RFC1: Reduced Folate Carrier 1
RIPA: Radioimmunoprecipitation assay
RPE: retinal pigmented epithelium
siRNA: short interfering RNA
SLC19a1: solute carrier family 19 (folate transporter), member 1
SNPs: single nucleotide polymorphisms
STING: stimulator of interferon genes
UV: ultraviolet
ZO-1: Zonula occludens 1

## Data availability

All data generated or analyzed during this study are included in this published article. Raw data and analyses can be provided upon request. There are no restrictions on data availability.

## Authors’ contributions

GG contributed to the study design, performed surgeries and experiments, obtained tissues, performed immunohistochemistry, collected images and data, analyzed the data, prepared figures, wrote the draft of the manuscript and contributed to editing. NB contributed to the design and execution of Western Blottings and collected membrane images. MCB designed and synthesized the lentiviral vector and performed cell-line Western Blottings. MSB contributed to the design and execution of qRT-PCR experiments and performed related analysis. GU contributed to the in vivo application of retinal ischemia model, retinal trypsin digestion and intravitreal injections, and writing of the paper. EK provided lentiviral vector as a courtesy and contributed to editing. MY acquired funds and administered the project, coordinated, and supervised the work, wrote the manuscript, and contributed to figures and edited the paper. All authors read and approved the final manuscript and provided minor modifications.

## Acknowledgments

We thank Dr. Canan Cakir Aktas (The Institute of Neurological Sciences and Psychiatry, Hacettepe University, Ankara, Turkey) for assisting the financial management of the project, Dr. Buket Nebiye Demir (The Institute of Neurological Sciences and Psychiatry, Hacettepe University, Ankara, Turkey) for technical advice on in vivo procedures, Dr. Cetin Demir (Faculty of Medicine, Department of Pediatrics, Division of Pediatric Oncology, Drug Resistance Laboratory, Hacettepe University, Ankara, Turkey) for his assistance in the execution of qRT-PCR experiments and our laboratory technician Mesut Firat (The Institute of Neurological Sciences and Psychiatry, Hacettepe University, Ankara, Turkey) for providing technical assistance throughout project. We are grateful to our undergraduate researchers Derin Nalcakan and Neslihan Nisa Gecici (Hacettepe University Medical School), who were officially added as interns to the project with “TÜBİTAK-Intern Researcher Scholarship Programme”, for their contribution to immunohistochemistry and data analysis.

**Supplementary information** is available for this paper.

**Correspondence and requests for materials** should be addressed to MY or GG.

